# Lack of circadian entrainment and limited nocturnal plasticity in response to a cyclic aversive stimulus in a diurnal rodent, the antelope ground squirrel (*Ammospermophilus leucurus*)

**DOI:** 10.1101/2024.09.27.615535

**Authors:** Victor Y. Zhang, G.J. Kenagy, Horacio O. de la Iglesia

## Abstract

Recent studies have shown that cyclic aversive stimuli, such as random footshocks, act as a nonphotic zeitgeber to entrain circadian behaviors in nocturnal rodents, a pattern termed “fear entrainment”. However, it remains unknown whether diurnal species exhibit similar plasticity in behavioral timing. This study aimed to determine if antelope ground squirrels (*Ammospermophilus leucurus;* AGS), a naturally diurnal rodent, can also shift their activity patterns to cyclic aversive stimuli. We conducted two experiments with 20 AGS housed in custom cages featuring a safe nesting area and a separate foraging area (for feeding and drinking), rendered aversive by presentation of unsignaled, time-specific footshocks. In the first experiment, animals were subjected to a 12:12 light-dark (LD) cycle. One group experienced aversion during the light phase, while the control group received the same treatment during the dark phase. In the second experiment, a 16:8 LD cycle was used, and animals were divided into three groups with the foraging area rendered aversive either during the first or second half of the light phase or during the dark phase. Following each of these treatments, animals were released into constant darkness (DD) to assess the phase and period of free-running rhythms. Contrary to previous findings in nocturnal rodents, AGS did not exhibit consistent shifts in activity to avoid footshocks. Most animals maintained their normal diurnal activity, with only minor and inconsistent phase shifts. In experiment two, animals exposed to footshocks during half of the light phase also failed to reliably shift activity to the opposite “safe” portion of the light phase. Together, these findings show AGS lack the ability to entrain to cyclic aversive stimuli or become nocturnal; this result suggests a lack of substantial plasticity in activity timing. These results highlight the importance of considering species-specific differences in nonphotic circadian entrainment and temporal niche plasticity.

## Introduction

To align routine behaviors and their physiological coordination with cyclic changes in the external environment, animals have evolved endogenous circadian clocks that generate and sustain rhythms with a periodicity of ∼24 h. The natural daily light-dark (LD) cycle is the principal environmental cue (or zeitgeber) that entrains the endogenous central circadian clock and its associated rhythmic processes in mammals (DeCoursey, 1972; Pittendrigh & Minis, 1964; Shigeyoshi et al., 1997). However, nonphotic cyclic cues such as food availability, temperature cycles, and social interactions can also entrain circadian rhythms (Castillo-Ruiz et al., 2012; Hut et al., 1999; Marchant et al., 1997; Menaker & Eskin, 1966; Mistlberger, 2011; Mrosovsky, 1988). This entrainment process may operate through molecular mechanisms and neural pathways that differ in their relationship from those by which light entrains the central pacemaker in the brain, the suprachiasmatic nucleus (SCN) (Johnson et al., 2003; Marchant et al., 1997; Marchant & Mistlberger, 1997; Stephan et al., 1979a, 1979b). Compared to photic entrainment, our understanding of entrainment by non-photic zeitgebers is limited.

Recent evidence has shown that time-specific aversive stimuli may act as a zeitgeber (Bussi et al., 2024; Pellman et al., 2015). Normally nocturnally active mice and rats placed in a specially designed cage with a safe nest and a separate chamber where they can retrieve all their food and water will shift their activity rhythms to the light phase when they are exposed to random electric footshocks during the dark phase. The diurnal rhythm persists under free-running conditions, in the absence of light cues and footshocks, and is also reliant on an intact amygdala and functioning molecular circadian clock within the SCN (Bussi et al., 2024). The adaptive value of entraining activity to avoid aversive stimuli (termed “fear entrainment”) could be to enhance survival by bolstering an individual’s ability to evade predators. Indeed, several studies conducted on free-living animals suggest that cyclic predator pressures can alter behavioral rhythms in prey species (Fenn & Macdonald, 1995; Ross et al., 2013; Sipari et al., 2016; Swarts et al., 2009; Vinne et al., 2019). All studies published so far examining potential shifts in activity timing in response to aversive stimuli have been conducted on nocturnal species, leaving open the question as to whether entrainment by aversive stimuli would occur in diurnal species.

The antelope ground squirrel (AGS) is a useful rodent model for investigating biological timekeeping in diurnal species because AGS remain exclusively diurnal when brought from the field into the laboratory (Kenagy, 1978; Refinetti & Kenagy, 2018a). They display robust, high-amplitude, daytime activity rhythms (Kenagy, 1978; Refinetti & Kenagy, 2018a) and possess an intrinsic circadian period that remains stable across the year (Kenagy, 1978). Notably, the similarity of the circadian period of AGS (24.2 h) (Kenagy, 1978; Refinetti & Kenagy, 2018b) to other diurnal mammals (including humans) (Aschoff, 1979) suggests similar mechanisms of entrainment compared to those of nocturnal species, which typically have periods shorter than 24 h (Johnson et al., 2003; Kenagy, 1978). Exploring the potential entraining effects of a cyclic aversive stimulus such as electric footshocks (and a potential resulting emotional response such as fear) in AGS could also provide insights into the kinds of circadian disruptions that have been widely observed in human anxiety disorders such as post-traumatic stress disorder (PTSD) (Lewis et al., 2009; Ohayon & Shapiro, 2000; Spoormaker & Montgomery, 2008).

In the present study with the AGS, we used a “fear cage system” based on the design by Pellman et al. (2015). We hypothesized that time-specific aversion to electric shocks could act as a zeitgeber to entrain circadian rhythms in a strictly diurnal rodent. Because previous research has shown that rats and mice presented with nocturnal footshocks shift rhythms of foraging and feeding to the light phase, we predicted that AGS presented with footshocks during the light phase would likewise shift the same behavioral functions to become nocturnal. Additionally, given the possibility that AGS may be limited in the plasticity of their activity timing, we also explored whether shorter durations of footshocks presented only during portions of the light phase would be sufficient to shift AGS rhythms to different times of the light phase. Specifically, we predicted that morning footshocks would entrain behaviors to the afternoon, while afternoon footshocks would entrain behaviors to the morning.

## Methods

### Animal collection and housing

Twenty antelope ground squirrels (6 females, 14 males) were captured in their natural desert habitat in Harney County, Oregon, in April 2021 using Sherman live traps (H. B. Sherman Traps, Inc. Tallahassee, Florida). Animals were transported to a laboratory facility at the University of Washington. Animals were housed in plastic cages (26cm wide × 47.6cm long × 15.2cm deep) with running wheels (17.5 × 7.5cm diameter × width), provided with corn cob bedding, and maintained under a 12:12h LD cycle (∼1500 lux:red light ∼1 lux, measured from middle of cage interior). Animals received an ad libitum diet of (PicoLab Rodent Diet 20 LabDiet 5053) and water, and were held in these general conditions for two months of adjustment to captivity and laboratory conditions before the experiments were initiated. The animals gained weight and maintained good health through the study. All studies were approved by the University of Washington Office of Animal Welfare and Institutional Animal Care and Use Committee (4045-02). Squirrel captures were conducted under the authorization of a Scientific Taking Permit (032-21) with the Oregon Department of Fish and Wildlife.

### Experimental chamber

Our cage design is based, with minor modifications, on the fear paradigm developed by Pellman et al. (2015). Our cages measured 26cm wide × 47.6cm long × 15.2cm deep, divided into two equal compartments by a black acrylic wall that contained one isolated experimental animal setup on each side, resulting effectively in two independent cages. The space within each individual unit was further divided into two halves consisting of an enclosed nest box and an open activity area with a wire grid floor, which was the only compartment where food and water were available and will be referred to as the “foraging chamber” hereafter. The nest chamber was an acrylic box (20.6cm wide × 10.5cm long × 18.1cm deep) with a 3.8cm-diameter hole that allowed access to the foraging chamber. A rectangular panel of 10.5 × 18.1cm on one side of the nest chamber was constructed of clear, rather than black, acrylic to allow daily inspections of the animal without additional disturbance. The nest chamber floor was provided with an 8-cm depth of corn cob bedding. The floor grid of the foraging chamber was composed of 16 stainless steel rods (4.5mm diameter) with spaces of 3.5-mm between the bars, which were connected to a precision animal shocker (Coulbourn Instruments, Allentown, PA). The overall dimensions of the foraging chamber floor were 12 × 21 cm; a water bottle was situated at one end and a food dispenser at the other.

Each cage chamber was equipped with three movement-detection sensors located as follows: (1) inside the lid of the nesting chamber, (2) above the center of the foraging chamber, and (3) within the opening of a food dispenser mounted outside of the chamber. Thus, we obtained a continuous record of activity associated with (1) movement of the animal within the nest box (nest activity), (2) movement of the animal in the foraging chamber (referred to as foraging activity hereafter), and (3) nose-pokes of the animal at the food dispenser. The sensors at (1) and (2) were passive infrared (IR) motion sensors (Product #189, Adafruit Industries, New York, New York), and the sensor (3) at the food dispenser was an IR break-beam sensor (Product #2167, Adafruit Industries, New York, New York). All sensor data were acquired and stored in 1-min bins on a computer using Clocklab software and hardware (Actimetrics, Wilmette, IL).

### Experimental design

Using the same individuals, we conducted two successive experiments, employing a different LD cycle and foot-shock schedule in each experiment. Before the first experiment, we allowed 14 days for animals to become accustomed to the new chamber under a 12:12 LD cycle. We then recorded 14 days of baseline activity from each animal. Following this, we randomly divided animals into two groups: light shock (LS) (N=12; 8 males and 4 females) and dark shock (DS) (N=8; 6 males and 2 females). LS animals were exposed to a 12-h interval of random footshocks during the light phase of the daily LD cycle, and DS animals were exposed to the same 12-h treatment during the dark phase. The footshocks consisted of activating the shock grid at 0.4mA for 10s at one random time within every 20-min interval. Following 15 days of these shock treatments, animals were released into constant darkness (DD) with no further shocks for 14 days to determine the free-running phase and period of their circadian rhythms.

Following the determination of free-running periods in DD of the first experiment, we began the second experiment, in which the squirrels were exposed to a 16:8 LD cycle. After 30 days of adjustment to the new photoperiod, we recorded baseline activity patterns for 14 days. Animals were then assigned to three groups: morning daytime shock (MS) (N=12; 10 males and 2 females), afternoon daytime shock (AS) (10 males and 2 females), and nighttime dark shock (DS) (N=4; 2 males and 2 females). All animals were exposed to an 8-h interval of random foot-shocks of 0.8mA intensity (twice the intensity we used in experiment 1) for 10s at one random time within every 20 min interval. The MS animals were presented with shocks during the first 8 hours of the 16-h light phase, the AS animals during the second 8 hours of the 16-hour light phase, and DS animals during the 8-h dark phase. The daily shock treatments of experiment 2 lasted for 15 days, after which animals were again released into constant darkness (DD) conditions for 20 days, with no further shocks. Because individual squirrels in the first experiment showed natural differences in baseline activity and chronotype (morning activity peak vs. afternoon activity peak vs. middle of day activity peak, or bimodal activity peaks), we balanced our assignment of individuals in the second experiment across treatment groups based on our visual determination of chronotype in experiment 1 to represent different chronotypes across groups.

### Analysis

All analyses were conducted with R statistical software (Version 4.3.3; http://www.R-project.org/) unless stated otherwise. Before conducting our analyses, we visually inspected the data and identified and removed occasional days of incomplete or fragmented periods of data collection or sensor failure.

To investigate changes in activity timing, we performed separate two-way mixed ANOVAs for each successive phase of each experiment (baseline, shocks, free-running) within the output of each sensor representing the different sensing areas of the experimental chamber (nest activity, foraging activity, and feeding). For each analysis, we used data from only the final four days of recordings for each phase and included the interaction between time (average 10min bins) and treatment (LS and DS in the first experiment and MS, AS, and DS in the second experiment) as fixed factors, including individual identity as a random effect.

We used recordings from the passive IR sensor located above the foraging chamber as our primary measure of circadian rhythms. This sensor captured the general movements of the animals within the foraging chamber, and these movements are likely less influenced by acute changes in feeding behavior due to shock treatment. Activity onsets were independently determined by four trained observers using the eye-fit method, and their estimates were averaged to calculate the mean projected phase of activity onset for each experimental phase and experiment. Using El Temps software (v.1.228, University of Barcelona, Spain), we also applied the cosinor method (Refinetti et al., 2007) to compute the phase of the fundamental component of foraging activity (acrophase) during the final four days of recordings from each experimental phase (Fig. S1). Rayleigh tests were then used to assess the average direction and magnitude of foraging activity for each treatment within each experiment, and confidence intervals were expressed as fiducial limits around the mean activity onset.

We used Watson-Williams tests to determine differences in mean phase between the different treatment groups at the time of release into DD. Actograms from individual recordings were also generated and used to visually assess the variable responses of individuals to shock treatment.

## Results

All animals entrained and showed diurnal activity patterns, as expected, to the initial LD cycles in both experiments 1 (12:12 LD) and 2 (16:8 LD). However, cyclic diurnal presentation of shocks failed to entrain any consistent nocturnal behavioral rhythms in AGS in either of our two experiments. AGS showed a wide range of inter-individual variation in responses to shock treatment, most of which was not suggestive of circadian entrainment to nocturnal activity or to any time of day when shocks were not being presented.

Surprisingly, we found significant interactions between treatment group and time across all cage sensors and experimental phases in the first experiment, suggesting that the timing of activity was dependent on treatment group assignment, even during the initial baseline period (Table 1a). For example, during the baseline period, LS animals appeared to have higher foraging activity than DS animals during the second half of the light phase (Fig. 1A). This interaction was even more pronounced during the shock presentation and free-running portions of experiment 1 (Fig. 1B, C). In contrast, during the second experiment, interactions between treatment and time were not observed during the initial baseline period, but rather only during the shock presentation and free-running portions of the experiment (Table 1b). For example, compared to DS and MS-treated animals, more nocturnal activity was seen in the foraging area of AS-treated animals during the shock presentation and free-running portions of experiment 2 (Fig. 1D-F).

**Table 1.** The influence of treatment grouping, time of day, and their interaction on activity measures in antelope ground squirrels during each experimental phase, as determined by two-way mixed ANOVAs. (a) Treatment groupings in experiment 1 include light shock (LS) and dark shock (DS), and (b) groupings in experiment 2 include morning shock (MS), afternoon shock (AS), and dark shock (DS).

| Variable | Experimental Phase | Treatment |  | Time |  | Interaction |  |
| --- | --- | --- | --- | --- | --- | --- | --- |
| a) Experiment 1 (12:12 LD) |  |  |  |  |  |  |  |
| Nest Activity | Baseline | F<sub>1,17</sub>=1.23 | P=0.28 | F<sub>143,10476</sub>=26.8 | P<0.0001 | F<sub>143,10476</sub>=2.50 | P<0.0001 |
|  | Shock | F<sub>1,17</sub>=8.51 | P=0.01 | F<sub>143,10207</sub>=31.2 | P<0.0001 | F<sub>143,10207</sub>=6.30 | P<0.0001 |
|  | Free-run | F<sub>1,17</sub>=0.71 | P=0.41 | F<sub>143,9775</sub>=31.7 | P<0.0001 | F<sub>143,9775</sub>=5.24 | P<0.0001 |
| Foraging Activity | Baseline | F<sub>1,18</sub>=0.03 | P=0.86 | F<sub>143,11194</sub>=19.7 | P<0.0001 | F<sub>143,11194</sub>=1.49 | P 0.0002 |
|  | Shock | F<sub>1,18</sub>=0.79 | P=0.39 | F<sub>143,10782</sub>=18.4 | P<0.0001 | F<sub>143,10782</sub>=9.33 | P<0.0001 |
|  | Free-run | F<sub>1,17</sub>=0.38 | P=0.55 | F<sub>143,10639</sub>=18.3 | P<0.0001 | F<sub>143,10639</sub>=6.29 | P<0.0001 |
| Feeding Activity | Baseline | F<sub>1,18</sub>=0.31 | P=0.59 | F<sub>143,11049</sub>=3.58 | P<0.0001 | F<sub>143,11049</sub>=1.93 | P<0.0001 |
|  | Shock | F<sub>1,18</sub>=4.82 | P=0.042 | F<sub>143,10345</sub>=4.06 | P<0.0001 | F<sub>143,10345</sub>=4.93 | P<0.0001 |
|  | Free-run | F<sub>1,17</sub>=0.03 | P=0.86 | F<sub>143,10639</sub>=2.99 | P<0.0001 | F<sub>143,10639</sub>=1.99 | P<0.0001 |
| b) Experiment 2 (16:8 LD) |  |  |  |  |  |  |  |
| Nest Activity | Baseline | F<sub>2,17</sub>=0.37 | P=0.69 | F<sub>143,10925</sub>=13.5 | P<0.0001 | F<sub>143,10925</sub>=0.81 | P=0.99 |
|  | Shock | F<sub>2,17</sub>=0.51 | P=0.61 | F<sub>143,10783</sub>=16.1 | P<0.0001 | F<sub>143,10783</sub>=3.44 | P<0.0001 |
|  | Free-run | F<sub>2,17</sub>=0.55 | P=0.59 | F<sub>143,11071</sub>=11.5 | P<0.0001 | F<sub>143,11071</sub>=3.46 | P<0.0001 |
| Foraging Activity | Baseline | F<sub>2,17</sub>=0.48 | P=0.63 | F<sub>143,10629</sub>=11.6 | P<0.0001 | F<sub>143,10629</sub>=0.94 | P=0.76 |
|  | Shock | F<sub>2,17</sub>=0.14 | P=0.87 | F<sub>143,11071</sub>=11.0 | P<0.0001 | F<sub>143,11071</sub>=1.61 | P<0.0001 |
|  | Free-run | F<sub>2,17</sub>=0.18 | P=0.84 | F<sub>143,10639</sub>=11.5 | P<0.0001 | F<sub>143,10639</sub>=1.26 | P=0.002 |
| Feeding Activity | Baseline | F<sub>2,17</sub>=0.08 | P=0.92 | F<sub>143,10223</sub>=2.97 | P<0.0001 | F<sub>143,10223</sub>=1.02 | P=0.38 |
|  | Shock | F<sub>2,16</sub>=0.95 | P=0.41 | F<sub>143,10496</sub>=1.36 | P=0.003 | F<sub>143,10496</sub>=1.37 | P<0.0001 |
|  | Free-run | F<sub>2,17</sub>=1.22 | P=0.32 | F<sub>143,11071</sub>=3.07 | P<0.0001 | F<sub>143,11071</sub>=3.08 | P<0.0001 |

**Figure 1:**
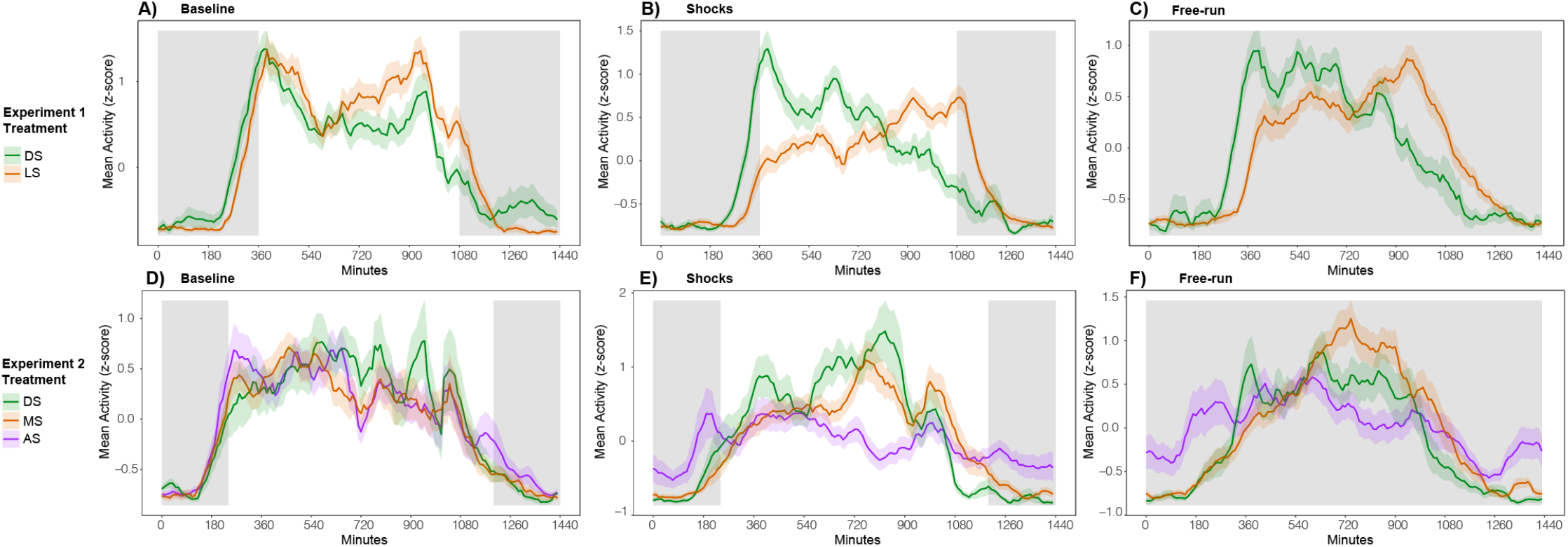
Average 24h waveforms of activity in the foraging area of across two experiments. Raw activity counts were normalized using a z-score and applying a 1h rolling mean. In the first experiment, squirrels were exposed to a 12:12 LD cycle and a 12h interval of shocks that were distributed either during the light or dark phase of the LD cycle (A-C). In the second experiment, squirrels were exposed to a 16:8 LD cycle and shocks were distributed during an 8h interval in the first half (“morning”) or the second half (“afternoon”) of the light phase, or in the dark phase. (D-F).

In the first experiment, Rayleigh tests for onsets (Fig. 2) indicate significant phase clustering of foraging activity for all experimental phases, suggesting that animals within each group coincided in the specific times at which they started becoming active. The projected phase of activity onset after transfer to DD differed significantly between DS and LS animals (Watson-Williams F1,18 = 7.30; p = 0.0146). However, the direction of this shift— resulting in a delay of approximately 2h in LS animals—did not help in avoiding shock exposure; instead, it had the opposite effect, shifting the animals’ activity further into the shock window (Fig. 2). In the second experiment, significant phase clustering of foraging activity onset (Fig. 2) was detected across all groups except in the DS animals during the initial baseline period and shock presentation phase, and there was no significant difference in projected phase of activity onset after transfer to DD (Watson-Williams F2,17 = 1.70; p = 0.21). Similar results were observed across both experiments when acrophase was used as the phase marker of activity (Fig. S1) Individual actograms showed that squirrels consistently remained diurnal during nocturnal shock presentations, but exhibited a wide variety of responses when shocks were presented during the light phase (Fig. 3). In response to shock presentation, squirrel behavioral rhythms showed either no response, a disruption, masking (acute avoidance of the shock intervals), or entrainment (Supplementary files “actograms”). Most phase shifts only enabled partial avoidance of footshocks over a two-week period, and entrainment to shock presentation (a complete shift of activity phase to avoid shocks) only occurred in two animals in experiment 2 (Fig. 3; bottom).

**Figure 2:**
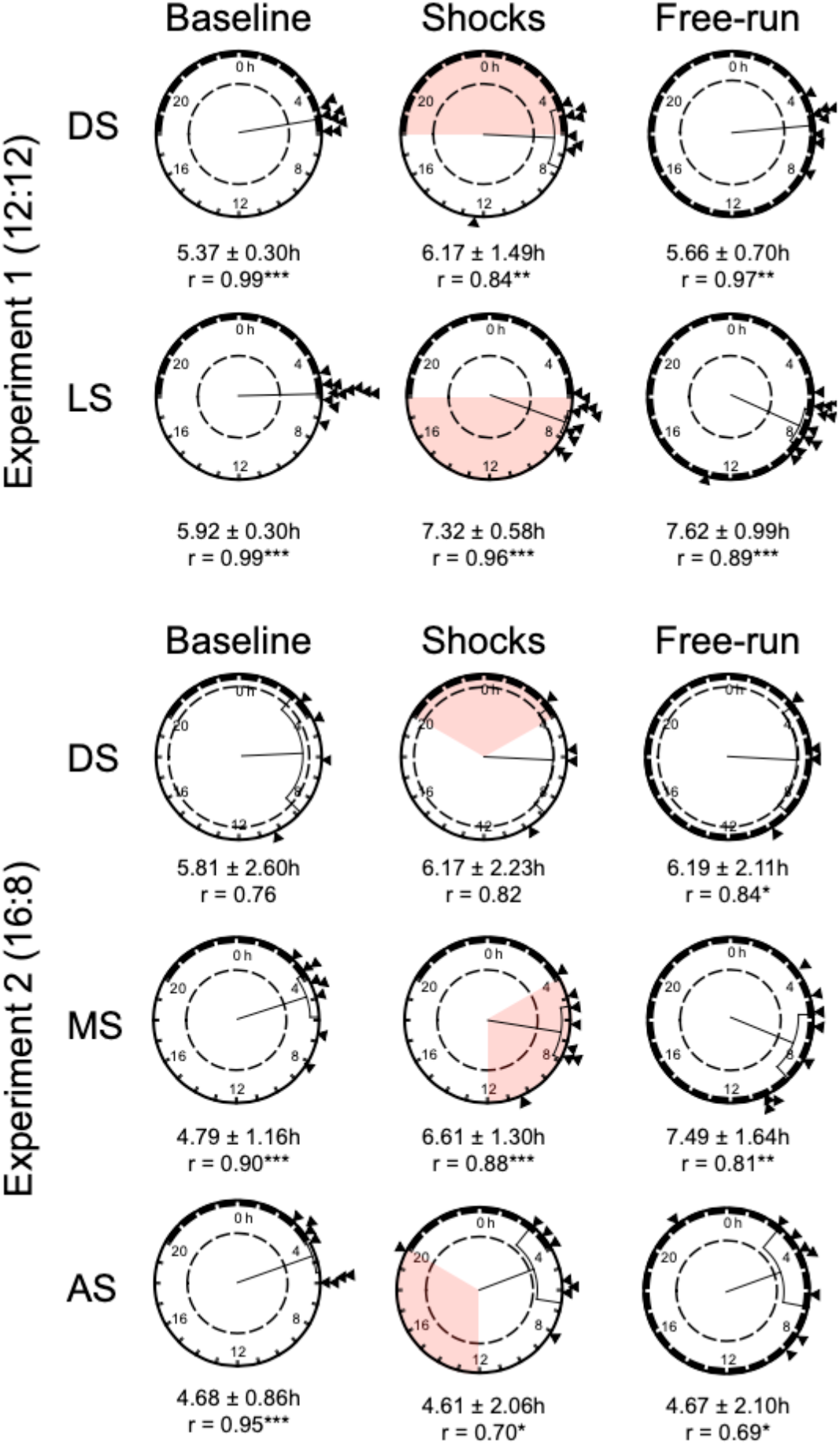
Circular 24-h plots showing direction and magnitude of mean foraging activity onset (shown as vectors) ± confidence intervals across two experiments. (Top) In the first experiment, squirrels were exposed to a 12:12 LD cycle and a 12h interval of shocks that were distributed either during the light or dark phase of the LD cycle. (Bottom) In the second experiment, squirrels were exposed to a 16:8 LD cycle and shocks were distributed during an 8h interval in the first half (“morning”) or the second half (“afternoon”) of the light phase, or in the dark phase. Red shading indicates interval of shock treatment. Rayleigh tests showed significant clustering of activity phase in all groups of experiment one, but not experiment two *\*p*<0.05m *\*\*p*<0.01, *\*\*\*p*<0.001.

**Figure 3:**
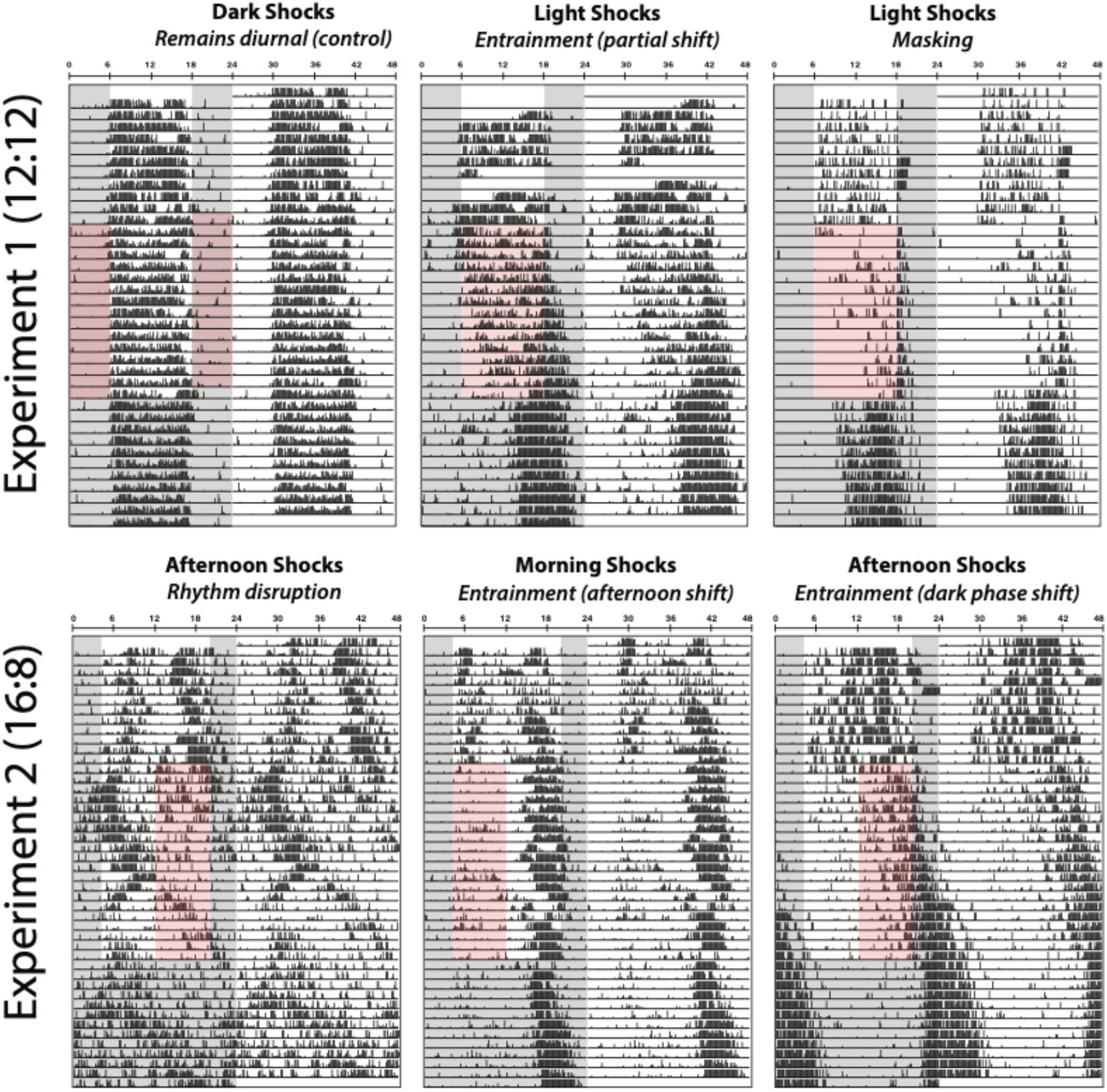
Double-plotted actograms of foraging activity in the experimental cage of six antelope ground squirrels from two experiments, shown to illustrate the diversity of behavioral responses to shock presentation. (Top) In the first experiment, squirrels were exposed to a 12:12 LD cycle and a 12h interval of shocks that were distributed either during the light or dark phase of the LD cycle. (Bottom) In the second experiment, squirrels were exposed to a 16:8 LD cycle and shocks were distributed during an 8h interval in the first half (“morning”) or the second half (“afternoon”) of the light phase, or in the dark phase. Gray shading indicates dark phase of the LD cycle and red shading indicates interval of random shocks.

## DISCUSSION

Most animal species synchronize their typical daily activity patterns to either the daytime or nighttime segments of the 24-h environmental cycle, along with which they show a suite of adaptive sensory modalities that are matched to either the nocturnal or the diurnal temporal niche. However, there is potential adaptive value to showing flexibility, which could allow a species to respond to an unusual environmental challenge. The plasticity of the temporal program of any species should be key to determining its ability to respond to such environmental changes. In this study we investigated whether cyclic aversive stimuli (footshocks) could serve as a zeitgeber to entrain circadian rhythms in a diurnal rodent, the AGS, by shifting from its normal diurnal activity to a nocturnal phase. We found that AGS did not exhibit entrainment to the aversive stimuli, contrasting with the recently well documented responses of nocturnal rats and mice (Bussi et al., 2024; Pellman et al., 2015) which shifted their activity to daytime when they received footshocks during the nocturnal phase of the 24-h cycle.

Mouse and rat studies alone are insufficient for providing a comprehensive understanding of how cyclic fear translates into rhythmic outputs, as the circadian neural circuitry of nocturnal species is not merely a diametrically opposed arrangement to that of diurnal species (Hut et al., 2012; Novak et al., 2008; Smale et al., 2008; Wams et al., 2017). While photic entrainment operates similarly in diurnal and nocturnal species (Daan & Pittendrigh, 1976; DeCoursey, 1972; Johnson et al., 2003), the responsiveness of individuals to entrainment by nonphotic zeitgebers can vary greatly across species and depend on the specific zeitgeber being considered and the neurosensory pathway by which it is perceived. For instance, time-restricted food availability strongly influences circadian phase preferences (López-Olmeda et al., 2010; Mistlberger, 2009; Stephan et al., 1979b), while induced locomotor activity (Hut et al., 1999; Reebs & Mrosovsky, 1989) and ambient temperature (Pohl, 1998; Refinetti, 2010; van der Vinne et al., 2014) play only a supplemental or reinforcing role to photic entrainment. Thus, investigating the effects of time-specific fear, as recently-characterized to serve as a nonphotic zeitgeber, for diurnal mammals is essential to gauge the generalizability of fear across prey species as a whole and to identify any potential variations in response patterns compared to nocturnal mammals.

For the few AGS individuals that exhibited changes in activity timing in response to shock treatment, phase shifts occurred at a notably slow rate, raising questions about the potential functional utility of this minor induced behavioral plasticity. For example, when exposed to the same electric shock paradigm, mice and rats shifted the timing of circadian activity onset by about 12 hours within 4-5 days (Bussi et al., 2024; Pellman et al., 2015). In contrast, all AGS, except one, failed to shift the timing of activity to avoid footshocks over a 2-3 week treatment period, exhibiting a pattern that reflected more their free-running rhythms than any semblance of entrainment by cyclic electric shock. The lack of entrainment by a cyclic aversive stimulus in AGS could be interpreted as a lack of perceived relevance of the cyclic stimulus. In other words, it is conceivable that the footshocks we administered were not strong enough to register an aversion by AGS and that the animals might not encode the stimulus as aversive. We do not believe that the animals did not perceive the shocks, because we visually confirmed that the animals reacted to the shocks by immediately retreating to their nest chambers, and most actograms reflect the active avoidance of the foraging chamber at, and immediately after, the moments when the shocks were presented.

Switches in daily activity timing, known as “temporal niche switching,” have been observed in free-living species in response to a variety of nonphotic environmental cues, including examples of predator pressures that drive such shifts (Fenn & Macdonald, 1995; Ross et al., 2013; Sipari et al., 2016; Swarts et al., 2009; Vinne et al., 2019). For instance, nocturnal rats shifted to diurnal activity when exposed to foxes that were nocturnally active (Fenn & Macdonald, 1995), and crepuscular island foxes (Zhang et al., 2022) reduced their daytime activity during periods of intense presence of diurnal golden eagles, but then increased daytime activity once the eagles were removed from their habitat (Hudgens & Garcelon, 2011; Swarts et al., 2009). However, to our knowledge, no studies have documented similar predator-induced temporal shifts in diurnal species. It seems that diurnal animals in general may be less likely to engage in temporal niche switching than their nocturnal and crepuscular counterparts (Hut et al., 2012). Several factors may be limiting diurnal species from engaging in such behavior, including their reliance on the visual system for orientation, foraging, and evading predators, and these sensory limitations may have led to evolutionary reductions in phenotypic plasticity (Gerkema et al., 2013; Lande, 2009). Because AGS possess a pure-cone retina (Crescitelli, 1961), they may lack the sensory capacity required to utilize the nocturnal niche. Additionally, phylogenetic inertia may have driven the majority of subfamilies within the family *Sciuridae* to become locked into the diurnal niche with little phenotypic plasticity. Our second experiment, which showed that AGS also failed to reliably shift activity to a different time within the light phase (morning or afternoon in a 16:8 LD cycle), supports this idea of phylogenetically constrained inflexibility. Specifically, even when cyclic aversive stimuli were presented during only part of the active light phase, AGS failed to shift their activity timing consistently to avoid electric shocks. Taken together, these observations suggest that AGS lack the plasticity to engage in significant nonphotic phase shifting regardless of illumination levels and at the intensity of shocks that we administered.

The absence of fear entrainment observed in this study may also be attributed to the absence of ecologically relevant cues that would drive temporal avoidance behaviors in squirrels. In the wild, the activation of defensive behaviors involves a trade-off between the benefits of immediate survival and the cost of feeding and reproduction, and thus should only be triggered when prey face a substantial risk of predation (Lima, 1998). Moreover, the induction of these behaviors depends on how information about danger is conveyed as well as the sensory mechanisms used by prey to detect potential threats. For instance, prey species utilize various direct signals for predator recognition, such as visual, acoustic, or olfactory cues. Indirect cues, like illumination levels, can also aid in avoidance behaviors (Orrock, 2004). In its natural environment, AGS are not active on the surface before dawn or after dusk (Germano & Saslaw, 2023), effectively excluding them from potential encounters with native crepuscular or nocturnal predators such as great horned owls (*Bubo virginianus*); during the day, AGS face a variety of predators, primarily diurnal raptors such as the red-tailed hawk (*Buteo jamaicensis*) and gopher snakes (*Pituophis catenifer*). Similar to other ground squirrel species, AGS likely rely on visual (Robinson, 1980) and auditory (Furrer & Manser, 2009) cues to respond to threats on the surface and olfactory cues to respond to threats in dark burrows where visual cues become unavailable (Coss & Owings, 1978; Hennessy & Owings, 1978). Given this, our experimental set up may have lacked appropriate ecological and evolutionary context to elicit natural fear responses that could potentially drive naturalistic behaviors of temporal avoidance.

Overall, the mechanisms underlying fear entrainment in nocturnal rodents (Bussi et al., 2024; Pellman et al., 2015) likely involve complex interactions between the SCN and extra-SCN oscillators. The molecular clock within the SCN is necessary but not sufficient for fear entrainment of circadian behaviors in mice, and the phase of clock gene expression within the SCN remains coupled to the LD cycle under conditions of fear-entrainment (Bussi et al., 2024). This suggests the existence of fear-entrainable oscillator(s) (FrEOs) that interact with the SCN to influence downstream behavioral outputs, though the specific locations of the FrEOs remain unidentified. Contrasting with findings from mice, our results suggest the possibility that some diurnal species either lack FrEOs or possess weak FrEOs that are easily overridden by the light-entrained SCN. This difference underscores the importance considering species-specific adaptations to circadian entrainment and corroborates existing literature showing that nocturnal and diurnal animals may possess fundamentally different mechanisms for entrainment by nonphotic zeitgebers (Novak et al., 2008; Wams et al., 2017).

Because the AGS is the first diurnal rodent species tested for the possibility of fear-entrainment, any generalizations about how the possibility of fear entrainment might function in diurnal mammals must be made cautiously. Considering the potential clinical significance of fear entrainment to human health, particularly in patients suffering from anxiety disorders such as PTSD, our results highlight the need for careful consideration when extrapolating findings from nocturnal rodent models to humans. The lack of entrainment observed in AGS may offer insights into the challenges of managing circadian and sleep disruptions associated with anxiety disorders in human patients. Understanding these differences will ultimately be essential for developing accurate models of how fear entrainment operates in mammals and its implications in both behavioral ecology and human health. Future research will also benefit from a comparative approach that includes ethologically relevant cues (e.g., predator-specific olfactory cues) to determine the relevance of fear-entrainment across mammalian species.

## Acknowledgements

We thank members of the de la Iglesia lab for their assistance in the lab.

## Author Contributions

VYZ, GJK, and HOD contributed to the development of the concept. VYZ and GJK captured animals in the field and provided husbandry in the lab. VYZ analyzed data, drafted figures, and wrote the initial manuscript with input from GJK and HOD. All authors contributed to revisions of the manuscript

## Statements and Declarations

### Ethical considerations

This study was conducted as per guidelines of the University of Washington International Animal Care and Use Committee (IACUC no. 4045-02). Work at our study sites and scientific taking were permitted by the Oregon Department of Fish and Wildlife (Oregon Permit no. 032-21) *Declaration of conflict of interest*: The author(s) declared no potential conflicts of interest with respect to the research, authorship, and/or publication of this article

### Data Availability

The datasets generated during and/or analyzed during the study are available from the corresponding author upon request.

## SUPPLEMENTARY FILES

**Figure S1:**
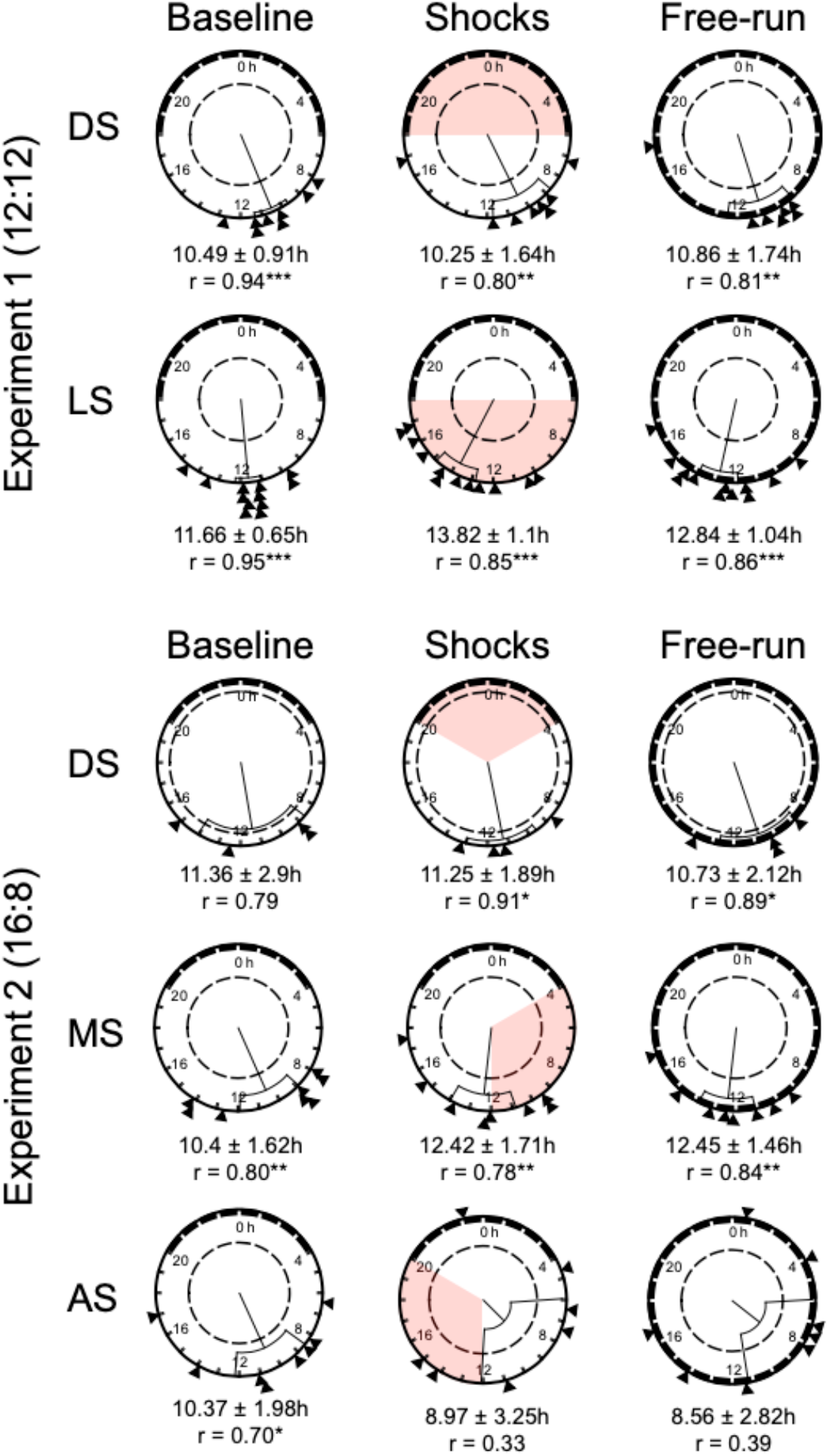
Circular 24-h plots showing direction and magnitude of mean foraging acrophase (shown as vectors) ± confidence intervals across two experiments. (Top) In the first experiment, squirrels were exposed to a 12:12 LD cycle and a 12h interval of shocks that were distributed either during the light or dark phase of the LD cycle. (Bottom) In the second experiment, squirrels were exposed to a 16:8 LD cycle and shocks were distributed during an 8h interval in the first half (“morning”) or the second half (“afternoon”) of the light phase, or in the dark phase. Red shading indicates interval of shock treatment. Rayleigh tests showed significant clustering of activity phase in all groups of experiment one, but not experiment two *\*p*<0.05m *\*\*p*<0.01, *\*\*\*p*<0.001.

